# Macroevolutionary dynamics of gene family gain and loss along multicellular eukaryotic lineages

**DOI:** 10.1101/2022.08.21.504376

**Authors:** Mirjana Domazet-Lošo, Tin Široki, Tomislav Domazet-Lošo

**Affiliations:** Department of Applied Computing, Faculty of Electrical Engineering and Computing, University of Zagreb, Unska 3, HR-10000 Zagreb, Croatia; Laboratory of Evolutionary Genetics, Ruđer Bošković Institute, Bijenička cesta 54, HR-10000 Zagreb, Croatia; School of Medicine, Catholic University of Croatia, Ilica 242, HR-10000 Zagreb, Croatia

## Abstract

The recent analyses in animals show that gene gain and loss fluctuate during evolutionary time. However, these studies lack the full profile of macroevolutionary trajectories as they do not consider other eukaryotic lineages and deeper evolutionary periods that go back to the origin of cellular organisms. To give a more inclusive view on the macroevolutionary patterns of genome complexity across the tree of life, we here recovered the evolutionary dynamics of gene family gain and loss ranging from the ancestor of cellular organisms to four focal species that represent animal, plant, and fungal diversity. We show that in all considered lineages the gene family content follows a common evolutionary pattern, where the number of gene families reaches the highest value at a major evolutionary and ecological transition, and then gradually decreases towards extant organisms. This pattern suggests that the genome complexity, approximated by the number of residing gene families, does not continuously increase through evolutionary time. We conclude that simplification by gene family loss is a dominant force in Phanerozoic genomes of various lineages, probably underpinned by intense ecological specializations and functional outsourcing. These findings support current theoretical predictions on the macroevolutionary change in genome complexity.

## Introduction

New genes are continuously added to genomes through evolutionary time (Domazet-Lošo and Tautz 2003, Tautz and Domazet-Lošo 2011). The mechanisms of their formation and their adaptive significance are widely discussed (Domazet-Lošo and Tautz 2003, Kaessmann 2010, Tautz and Domazet-Lošo 2011, Khalturin et al. 2009, Chen et al. 2013). However, the lifecycle of genes also includes gene loss (Neme and Tautz 2014), which is comparably much less studied process (Wolf and Koonin 2013, Albalat and Cañestro 2016, O’Malley et al. 2016). The recent work reveals that the reductive evolution by gene loss, in parallel with gene gain, played a major role in the evolution of animals (Richter et al. 2018, López-Escardó et al. 2019, Guijarro-Clarke et al. 2020, Fernández and Gabaldón 2020). However, these studies do not consider gain-and-loss patterns in other eukaryotic branches, and do not cover phylogenetic nodes deeper than the unicellular ancestors of animals (Richter et al. 2018, Guijarro-Clarke et al. 2020, Fernández and Gabaldón 2020) or the last common ancestor of fungi and animals (López-Escardó et al. 2019). The deepest node reached so far are the split between eukaryotic lineages Amorphea and Diaphoretickes (Bakarić 2016), but this early work is not directly comparable to later studies (Richter et al. 2018, López-Escardó et al. 2019, Guijarro-Clarke et al. 2020, Fernández and Gabaldón 2020), because it used a non-standard sequence similarity search algorithm with substantially less sensitivity compared to BLAST.

The macroevolutionary patterns of gene gain and loss are inseparable from the evolution of genome complexity — another long-standing question that in principle could be traced by reconstructing the content of ancestral genomes (Wolf and Koonin 2013, O’Malley et al. 2016). Previous work that aimed to recover ancestral genome content was largely restricted to specific phylogenetic nodes that represent, for example, eukaryotic (Rogozin et al. 2012), holozoan (Grau-Bové et al. 2017, Richter et al. 2018) and metazoan ancestor (Richter et al. 2018, Paps and Holland 2018, Fernández and Gabaldón 2020). In these attempts to estimate the ancestral genome complexity, researchers used various genome features like introns (Rogozin et al. 2012, Grau-Bové et al. 2017), orthologous genes (e.g., Grau-Bové et al. 2017, Richter et al. 2018, Paps and Holland 2018, Guijarro-Clarke et al. 2020, Fernández and Gabaldón 2020), and protein domains (e.g., Grau-Bové et al. 2017, López-Escardó et al. 2019). However, a comprehensive effort that would encompass the full macroevolutionary profiles from the origin of life to extant organisms is missing. The lack of such information largely precludes testing the proposed models of genome complexity evolution (Wolf and Koonin 2013, McShea and Hordijk 2013, O’Malley et al. 2016). According to the biphasic model, the episodes of rampant increase in genome complexity that are achieved by gene gain are followed by protracted periods of genome simplification through gene loss (Wolf and Koonin 2013). Similarly, complexity-by-subtraction model predicts initial rapid increase of complexity followed by decrease toward an optimum level over macroevolutionary time (McShea and Hordijk 2013). However, some authors do not agree with these views and suggest that interactions between simplification and complexification are not predictable (O’Malley et al. 2016).

The recovering of gene gain-and-loss patterns on deep phylogenies with hundreds of terminal taxa is further limited with the speed and sensitivity of sequence similarity search algorithms. The recent development of next generation sequence similarity tools (Hauser et al. 2013, Steinegger and Söding 2016, Steinegger and Söding 2017), which, by keeping comparable sensitivity, substantially outperform older solutions like BLAST in terms of speed, opens opportunity for recovering gene gain-and-loss patterns on the larger collection of genomes. Another uncertainty in reconstructing genomic gain-and-loss patters relates to the choice of evolutionary units that are tracked down. For instance, one could opt to follow evolutionary patterns of protein domains, orthologous genes, gene duplicates (paralogs), or gene families (homologs). The choice will largely depend on which evolutionary question one aims to address. So far, gain-and-loss studies mainly focused on tracing orthogroups (Paps and Holland 2018, Richter et al. 2018, Guijarro-Clarke et al. 2020, Fernández and Gabaldón 2020), presumably because it is assumed that orthologous genes are functionally more conserved than other homologs, although this view is repeatedly challenged (Demuth and Hahn 2009, Stamboulian et al. 2020). Nevertheless, we and others showed that significant shifts in the protein sequence space, which could be recovered by unrestrictedly tracing homologs (gene families), also carry the information on macroevolutionary important adaptive events (Domazet-Lošo et al. 2007, Domazet-Lošo and Tautz 2010a, Domazet-Lošo and Tauz 2010b, Quint et al. 2012, Domazet-Lošo et al. 2017, Trigos et al. 2017, Shi et al. 2020, Futo et al. 2021).

With an aim of getting a broader perspective on gene family gain-and-loss patterns and to test genome complexity dynamics across the tree of life, we here analyzed evolutionary lineages that start at the common ancestor of cellular organisms and end up in four focal species that represent the diversity of deuterostomic animals (*Homo sapiens*), protostomic animals (*Drosophila melanogaster*), plants (*Arabidopsis thaliana*), and fungi (*Saccharomyces cerevisiae*).

## Results

### Gene family content along four lineages

To cover all nodes on the consensus phylogeny that embraces our four focal species (Supplementary File 1), we retrieved 667 reference genomes (see Methods). This number of reference genomes is approximately two-fold higher compared to previous studies (Richter et al. 2018, López-Escardó et al. 2019, Guijarro-Clarke et al. 2020, Fernández and Gabaldón 2020). We then clustered the amino acid sequences of these reference genomes using MMseq2, a clustering tool which offers a fast and sensitive solution for large protein datasets (Hauser et al. 2016, Steinegger and Söding 2017). In contrast to previous studies that considered the gain- and loss patterns of orthologous groups (Richter et al. 2018, Guijarro-Clarke et al. 2020, Fernández and Gabaldón 2020) or protein domains on phylogenies (López-Escardó et al. 2019), we here explicitly traced homologous groups (gene families) (Domazet-Lošo and Tautz 2003, Demuth and Hahn 2009). We took this perspective since we were interested in recovering significant macroevolutionary changes in the protein sequence space (Domazet-Lošo and Tautz 2003, Domazet-Lošo et al. 2007, Domazet-Lošo and Tautz 2010a, Domazet-Lošo and Tauz 2010b, Domazet-Lošo et al. 2017, Futo et al. 2021); i.e., we aimed to estimate the overall dynamics of gene family diversity on the tree of life.

In this type of analysis, ancient and ubiquitous domains tend to attract multidomain proteins in large clusters, which are then placed at deep phylogenetic nodes (Domazet-Lošo and Tauz 2010b). To account for this effect, we varied the c-value of MMseq2 clustering algorithm, which sets the minimal percentage of sequence alignment overlap between proteins in a cluster, in the range between 0 and 0.8 (see Methods). On one extreme, a c-value of 0.8 forces clustering with alignment overlap of at least 80% protein sequence length. In turn, the obtained clusters contain protein sequences with highly similar overall domain architecture. In contrast, a c-value of 0 allows clustering without restrictions on the alignment overlap length that consequently leads to the formation of larger clusters, whose members can share only relatively short homologous regions. Expectedly, we find that the total number of gene families in all focal species depends on c-values. When we set no restriction on the length of alignment overlap (c-value = 0), we recovered the lowest number of gene families in total (Supplementary Table 1). In contrast, when we set the alignment length overlap at 80% (c-value = 0.8), we recovered the highest number of gene families (Supplementary Table 1).

Depending on the taxonomic composition of proteins in a cluster, we determined the evolutionary emergence of that gene family and its eventual loss on the consensus phylogeny (Supplementary File 1). This procedure allowed us to determine a gain-and-loss pattern for every protein family (cluster) recovered in the analyses (see Methods). We then used this information to reconstruct the protein family content of ancestral genomes along the focal species lineages (Fig. 1, Supplementary Table 1). In all four focal species we recovered a common evolutionary pattern where the number of gene families per ancestral genome abruptly increases at the origin of eukaryotes, keeps high values in early eukaryotic lineages, and after reaching a turning point continuously decreases towards extant organisms (Fig. 1). The phylogenetic position of nodes (phylostrata – ps) with the highest number of protein families, depends to some extent on the choice of c-values. However, regardless of these shifts in the phylogenetic position of the maximal peaks, the overall increase-peak-decrease pattern is always preserved (Fig.1).

**Figure 1.**
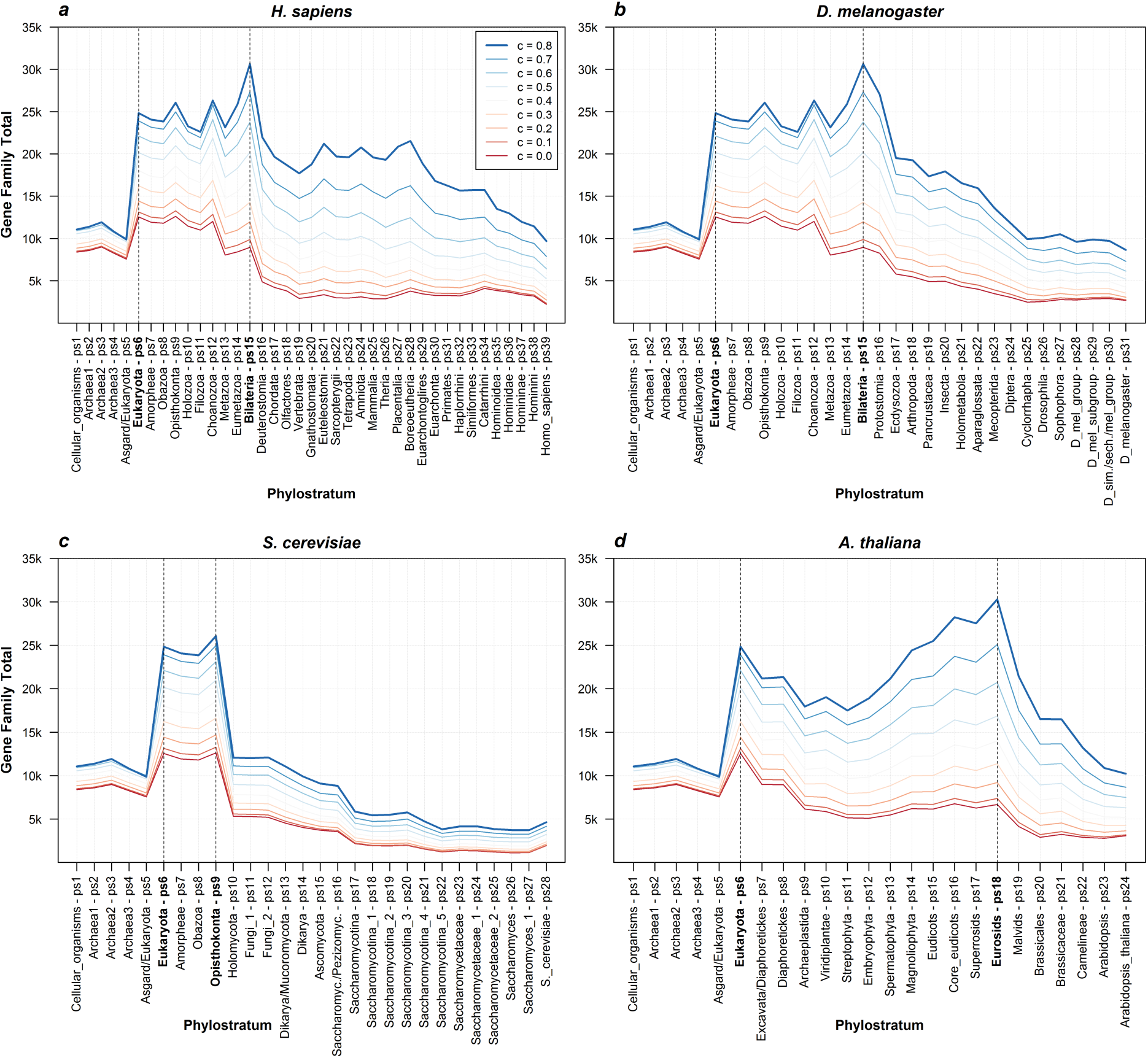
The total number of gene families along four focal lineages. The total number of estimated gene families across phylostrata (ps) is depicted for each of the four focal species (*H. sapiens*, *D. melanogaster*, *S. cerevisiae*, *A. thaliana*). The first phylostratum (ps1) represents the ancestor of cellular organisms and the last one corresponds to a focal species (e.g., ps39 for *H. sapiens*). Colored lines in each plot correspond to different c-values of the MMseq2 clustering algorithm. This parameter determines the minimal percentage of protein sequence alignment overlap in a cluster. The darkest blue graph corresponds to c-value = 0.8 which forces at least 80% of sequence length alignment overlap. The darkest red graph corresponds to c-value = 0 which allows clustering without restrictions on the alignment overlap length. The dashed vertical lines on each plot mark the evolutionary range with the highest number of gene families.

For instance, in the analyses of *H. sapiens* and *D. melanogaster* lineage (Fig. 1a,b), the maximum number of protein families varies between Eukaryota (c-value 0, ps9) and Bilateria (c-value 0.8, ps15). These patterns suggest that the early eukaryotes had the richest catalogue of protein families, if one takes the overall homology into account and ignores the precise architecture of multidomain proteins (c-value 0). In contrast, if within-gene synteny of homologous regions is tightly controlled by forcing high percentage of alignment overlap (c-value 0.8), then the first bilaterians show the richest collection of protein families. In either case, the number of protein families more or less continuously decreases after these maximal peaks (Fig. 1a,b). Regardless which criteria for gene family reconstruction we applied, these patterns suggest that ancestral genomes around the origin of animals and during their early diversification were more complex than the genomes of present animals in terms of gene family diversity.

In the analysis of *S. cerevisiae* lineage, we found the maximum number of protein families in the range between Eukaryota (ps6) and Opisthokonta (ps9) (Fig. 1c). The c-value profiles have very similar shape, which suggests that multi-level reductive processes dominated at the onset and during diversification of the fungal lineage (Holomycota - ps10 to *S. cerevisiae* - ps28, Fig. 1c). In contrast, the *A. thaliana* profile is more complex and uncovers that the ancestor of Eukaryota (ps6) had the highest diversity of gene families under permissive clustering (c-value 0). On the other hand, the stringent clustering, which forces within-gene synteny (c-value 0.8), reveals that the ancestor of Eurosids (ps18) harbored the highest number of gene families (Fig. 1d). However, all gene family profiles in the *A. thaliana* analysis revealed that the protein diversity loss is especially prominent after the origin of Eurosids (ps18) (Fig. 1d).

When compared to the suggested models of macroevolutionary change in genome complexity, our increase-peak-decrease patterns of gene family diversity fits the best the complexity-by-subtraction model, which envisages initial rapid increase in complexity that leads to a maximum value, followed by complexity decrease over macroevolutionary time towards an optimum (McShea and Hordijk 2013). However, our pattern also agrees with the biphasic model that predicts the episodes of rampant increase in genome complexity, which are followed by protracted periods of genome simplification (Wolf and Koonin 2013). The major difference between these two models is that the biphasic model predicts several waves of genome complexity change over macroevolutionary time (Wolf and Koonin 2013, McShea and Hordijk 2013). In our gene family analysis, we see an obvious two-wave gene family diversity pattern only in *A. thaliana* lineage where the genome complexity measured by gene family diversity first peaks at Eukaryota (ps6, Fig. 1d), decreases towards Streptophyta (ps11, Fig. 1d), peaks again at Eurosids (ps18, Fig. 1d), and then finally rapidly decreases towards focal *A. thaliana* (ps24, Fig. 1d).

It is important to note that the type of evolutionary information, conveyed by gene family profiles in Fig. 1, depends on the c-value. On one extreme, when clustering is performed with c-value = 0 the recovered gene families reflect the evolution of completely new sequences in the protein sequence space. On the other extreme, when clustering is performed with c-value = 0.8 the recovered gene families reflect the evolution of specific protein sequence architectures along the most of their sequence length. Accordingly, the highest number of gene families at Eukaryota (ps6, Fig. 1) in profiles generated with c-value = 0, suggests that evolution through the emergence of entirely new protein sequences were the most pronounced during eukaryogenesis. In contrast, the highest number of gene families in profiles generated with c-value = 0.8 at Bilateria (ps15, Fig. 1a,b) and at Eurosids (ps18, Fig. 1d) reveal<u>s</u> that the emergence of proteins with strictly defined protein architecture were at peak in these periods of animal and plant evolution. These results agree with a previous work in animals which detected an increased acquisition of new genes via protein sequence rearrangement processes at the stem of animals (López-Escardó et al. 2019). However, in the yeast analysis there are no apparent shifts in the phylogenetic position of maximal number of protein families when the c-values profiles of 0 and 0.8 were compared (Fig. 1c). This suggests that in the fungal lineage the emergence of completely new protein sequences and the assembly of proteins with specific sequence architectures were largely in balance.

### Gain and loss ratios

Although our gene family content profiles in Fig. 1 are the result of a careful bioinformatic analysis in a broad parameter space, the clustering of evolutionary distant reference genomes in large numbers and with new algorithms could possibly lead to unforeseen biases that obscure evolutionary signals and distort biological reality. We therefore made a more detailed evaluation of our gene family datasets, compared recovered patterns to findings in previous studies, and looked for expected and novel biological imprints. First, we explored the ratios of gene family gain and loss across lineages that lead to our four focal species (Fig. 2, Supplementary File 2). We found substantial loss events at the origin of metazoans (ps13) and deuterostomes (ps16) in the analysis that takes *H. sapiens* as a focal species (Fig. 2a). Similarly, on the lineage leading to *D. melanogaster*, we found extensive loss events at the origin of metazoans (ps13), protostomes (ps16), and ecdysozoans (ps17) (Fig. 2b). These results fully agree with previous studies which, by tracing orthologs groups, detected major loss events at the origin of metazoans (Richter et al. 2018, Guijarro-Clarke et al. 2020), deuterostomes (Fernández and Gabaldón 2020, Guijarro-Clarke et al. 2020), protostomes (Fernández and Gabaldón 2020, Guijarro-Clarke et al. 2020), and ecdysozoans (Fernández and Gabaldón 2020, Guijarro-Clarke et al. 2020).

**Figure 2.**
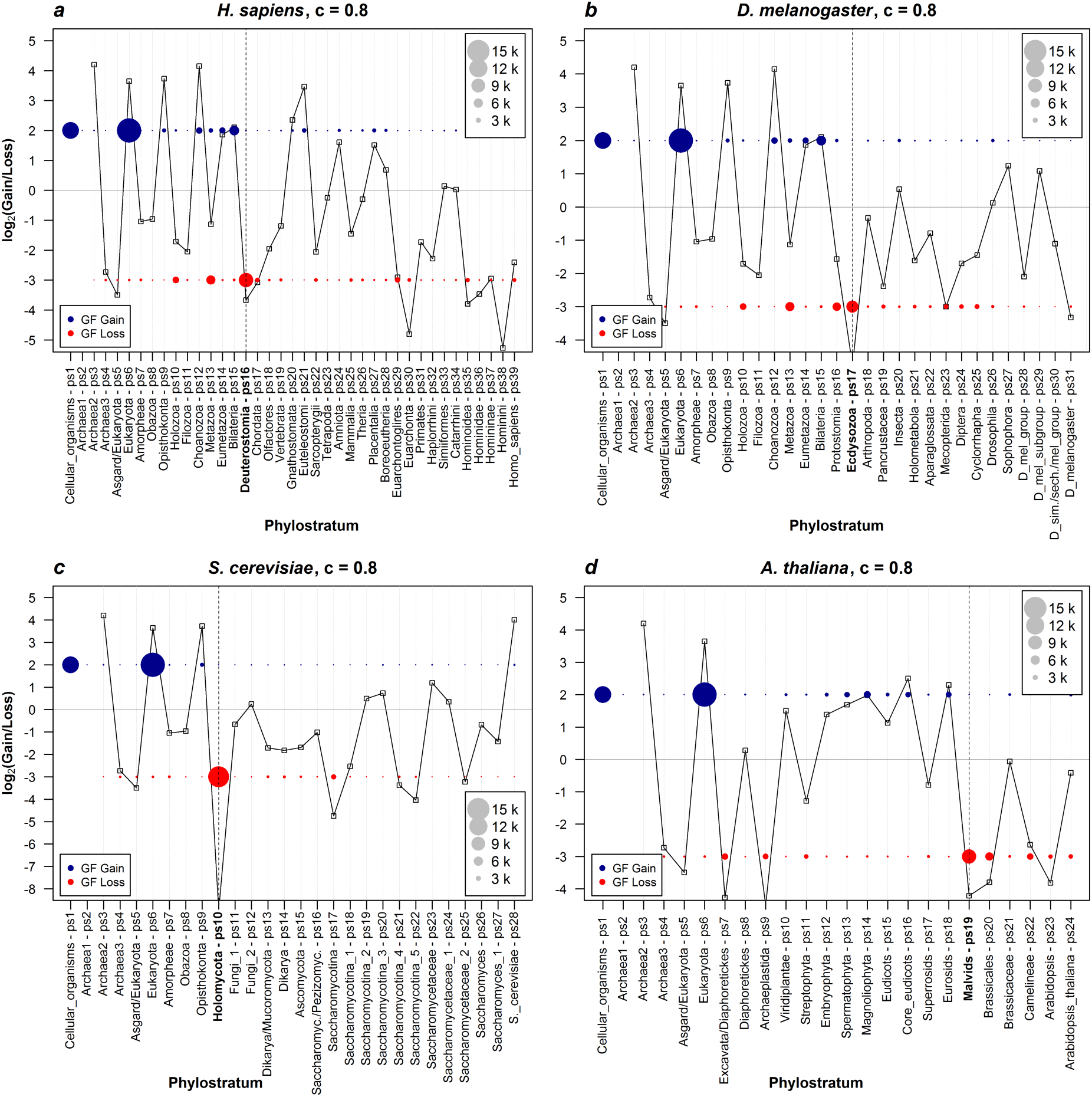
Evolutionary dynamics of gene family gain and loss ratios. The log_2_ values of the gene family gain and loss ratio across phylostrata for each of the four focal species (*H. sapiens, D. melanogaster, S. cerevisiae, A. thaliana*) are shown. We recovered gene families using the c-value cutoff of 0.8. Figures with other c-values cut-offs (0 to 0.8) are shown in Supplementary File 2. The first phylostratum (ps1) represents the ancestor of cellular organisms and the last one corresponds to a focal species (e.g., ps39 for *H. sapiens*). The vertical dotted lines mark the phylostrata with the major gene family loss events that precede the changing trends in the total number of protein families (Fig. 1, c-value 0.8). The circles of different sizes represent the number of gained (blue) and lost (red) gene families in a respective phylostratum.

The gain and loss profile in the *S. cereviseae* lineage reveals a dominant loss event at Holomycota (ps10) and a largely reductive trend in subsequent phylostrata (Fig 2c). This pattern agrees with previous work that detected substantial reductive genome evolution in fungi (Shen et al. 2018, Kiss et al. 2019). On the other hand, previous studies on gain and loss patterns in plants are rare and those that exist are not directly comparable to our study (e.g., Guo 2013, Bakarić 2016). However, the analysis of *A. thaliana* lineage shows an interesting two wave pattern (Fig. 1, Fig. 2d), where the first wave of dominant gene family losses stretches between Excavata/Diaphoretickes and Streptophyta (Fig. 2d, ps7 – ps11), and the second one between Malvids and Arabidopsis (Fig. 2d, ps19 – ps24). This result suggests that the diversification of land plants (Embryophyta to Eurosids, ps12 – ps18) was underpinned by substantial increase in gene family diversity (Fig. 1d, Fig. 2d).

### COG functional enrichments

We next sought to map functional data on our clusters and to explore if the obtained patterns are evolutionary meaningful. We first mapped COG functional categories to our phylogenetic pattern of gene family gain and loss, and then we explored enrichment profiles of every COG functional category across phylostrata (Fig. 3, Supplementary Files 3-6). In our four focal lineages we found that phylostrata, which show the enriched gain of some functional category generally precede the phylostrata that show the enriched loss of the same function (Fig 3, Supplementary Files 3-6). We also detected the two periods that are particularly enriched with functional gains. The first one is, conceivably, the origin of cellular organisms (ps1), which is enriched with the gains of essential metabolic functions, while the second one is the origin of eukaryotes (ps6), which is enriched with the gains of eukaryotic specific functions (Fig 3, Supplementary Files 3-6). Clusters without annotations show expected pattern with high gain and loss turnover in younger phylostrata (Fig 3, Supplementary Files 3-6, Tautz and Domazet-Lošo 2011, Palmieri et al. 2014). These global COG profiles reassured us that our clustering approach is evolutionary and biologically meaningful, hence we further refined the functional analysis by mapping fine grained Gene Ontology (GO) terms on our clusters.

**Figure 3.**
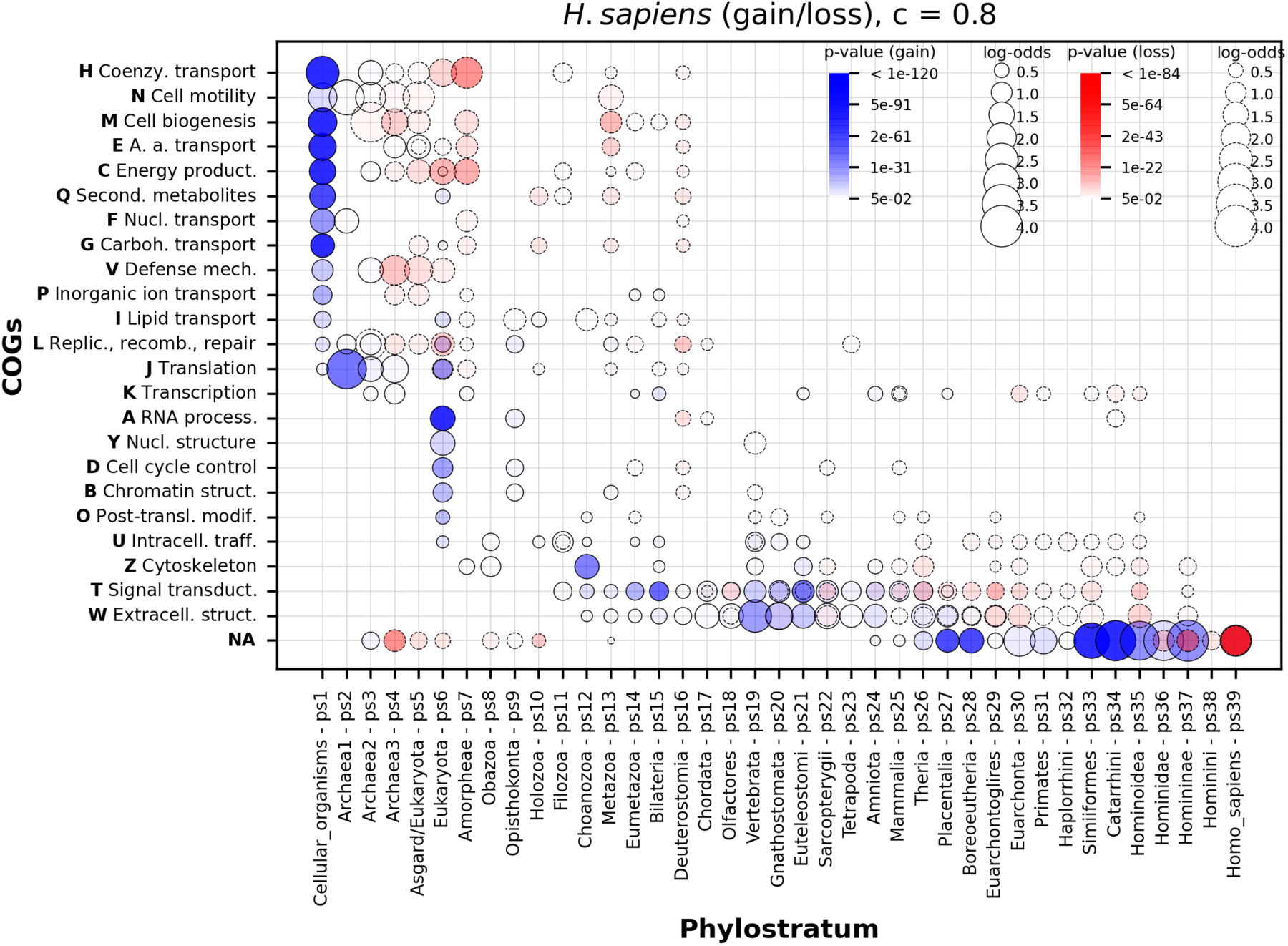
The enrichment of COG functional categories in gained and lost gene families along the *H. sapiens* lineage. The abbreviated names of COG functional categories and corresponding one-letter symbols are depicted at y-axis. The protein families without COG annotation are annotated with NA. The names and symbols of phylostrata are shown on the x-axis. The gene families are reconstructed with the MMSeq2 *cluster* program using a c-value of 0.8. Figures with c-values in the range between 0 and 0.8 for all four species are in Supplementary Files 3-6. Functional categories significantly enriched among gained gene families across phylostrata are depicted by solid circles painted in shades of blue that reflect p-values. The size of circles is proportional to enrichment values estimated by log-odds. Functional categories significantly enriched among lost gene families across phylostrata are depicted by dashed circles painted in shades of red that reflect p-values. The size of circles is proportional to enrichment values estimated by log-odds. The significance of enrichment was estimated by hypergeometric test corrected for multiple testing.

### GO functional enrichments

The full overview of profiles with enriched GO functions along our four lineages is depicted in Supplementary Files 7-10 (gain) and Supplementary Files 11-14 (loss). This large collection that contains approximately between 5,000 and 10,000 statistically significant GO enrichment profiles per focal species represents a rich reference resource for studying the evolution of gene family functions (Supplementary Files 7-14). In assessing these results, we first noticed that there are around two to three times more GO functional categories that are significantly enriched among gained compared to lost gene families (Table 1, Supplementary Table 2). However, because not all gene families have gene loss events − i.e., some gene families are retained up to a focal species − the number of functional enrichments in the gene family gain and loss pools are not directly comparable. To correct for this, we excluded from the analysis gene families that have only gain events (Table 1, Supplementary Table 2). In this way we were able to contrast the number of functional enrichments between gain and loss events using the same gene family pool. Regardless of these corrections, we again found the higher number of significantly enriched GO functions among gain events (Table 1, Supplementary Table 2).

**Table 1.** GO functions are more often enriched among gene family gain than loss events along four focal lineages.

| Number (%) of Enriched GO Functions |  |  |  |
| --- | --- | --- | --- |
| Focal species,<br>c-value | Gain | Loss | Gain/Loss Ratio |
| *H. sap.* c=0.8 | 6,407 (27.35) | 1,808 (8.1) | **3.38** |
| *D. mel.* c=0.8 | 6,247 (27.21) | 1,709 (7.79) | **3.49** |
| *S. cer.* c=0.8 | 2,895 (13.24) | 889 (4.33) | **3.06** |
| *A. thal.* c=0.8 | 4,179 (19.14) | 1,537 (7.71) | **2.48** |
| Number (%) of Enriched GO Functions (corrected) |  |  |  |
| Focal species,<br>c-value | Gain | Loss | Gain/Loss Ratio |
| *H. sap.* c=0.8 | 5,241 (23.47) | 1,808 (8.1) | **2.9** |
| *D. mel.* c=0.8 | 5,420 (24.69) | 1,709 (7.79) | **3.17** |
| *S. cer.* c=0.8 | 2,294 (11.17) | 889 (4.33) | **2.58** |
| *A. thal.* c=0.8 | 3,819 (19.15) | 1,537 (7.71) | **2.48** |

This robust asymmetry indicates that gene family gains are more often functionally non-randomly distributed over phylogeny compared to gene family losses. In other words, gene family gain events tend to functionally group together on the phylogeny, in contrast to gene family loss events which functions are more often randomly distributed and thus are not detected in the enrichment analysis. In the context of discussion on the genome complexity, this is an important finding because it supports the prediction of the biphasic model that genome simplification by gene loss is largely neutral process that occurs roughly in a clock-like manner (Wolf and Koonin 2013), in contrast to genome complexification by gene gain which usually occurs abruptly and is associated with evolutionary adaptations (Domazet-Lošo et al. 2007, Wolf and Koonin 2013).

After considering these global functional enrichment patterns, we sorted out some prominent functional categories, which demonstrate that our clustering approach and phylostratigraphic mapping recovers relevant biological information. First, we dissected the most striking increase in the number of gene families at the origin of eukaryotes (ps6 in all four lineages, Fig. 1,2). We recovered significant enrichment signals at the origin of eukaryotes (Fig. 4, Fig. 5, Supplementary File 15), related to gene family gains, for essentially all functions that are proposed in the literature to be eukaryotic defining properties (O’Malley et al. 2019, Cavalier-Smith et al. 2002, Koumandou et al. 2013). These include nucleus, endomembrane system, cytoskeleton and motility, endosymbiont, sexual reproduction and other eukaryotic specific features (Fig. 4, Fig. 5, Supplementary File 15). This result indicates that the abrupt increase in the number of protein families that we detected at Eukaryota (ps6, Fig.1,2) reflects the burst of innovations related to cell biology that occurred during eukaryogenesis (Koonin 2015, O’Malley et al. 2019, López-García and Moreira 2020).

**Figure 4.**
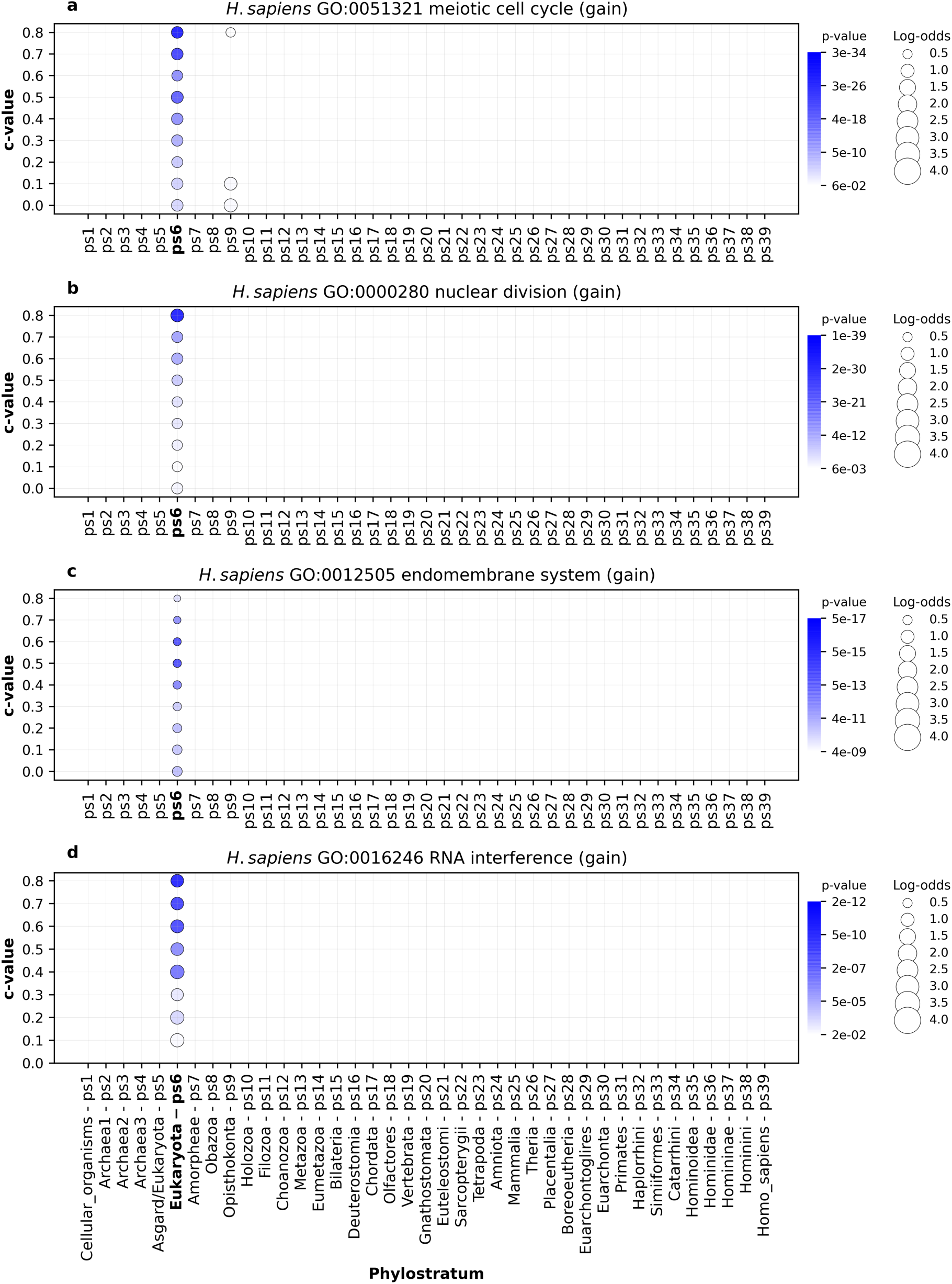
An example of GO functional enrichments related to eukaryogenesis in *H. sapiens* lineage. The results are shown for four GO terms: **a,** GO:0051321 (meiotic cell cycle) **b,** GO:0000280 (nuclear division) **c,** G:0012505 (endomembrane system) **d,** GO:0016246 (RNA interference). Functional enrichments were calculated using the set of gained gene families along *H. sapiens* lineage (x-axis). The extended catalog of eukaryogenesis-related terms is listed in Fig. 5 and their corresponding enrichment charts are shown in Supplementary File 15. The gene families are reconstructed with MMSeq2 *cluster* using a range of c-values (0 to 0.8, y-axis). Solid circles depict significant enrichments of a GO term in gene families gained at a particular phylostratum. The size of circles is proportional to an enrichment value estimated by log-odds, while the shades of blue correspond to p-values. The significance of enrichments was estimated by hypergeometric test corrected for multiple comparisons. Only enrichments with p < 0.05 are shown. The four depicted GO terms showed a consistent enrichment at Eukaryota (ps6) across tested c-values.

**Figure 5.**
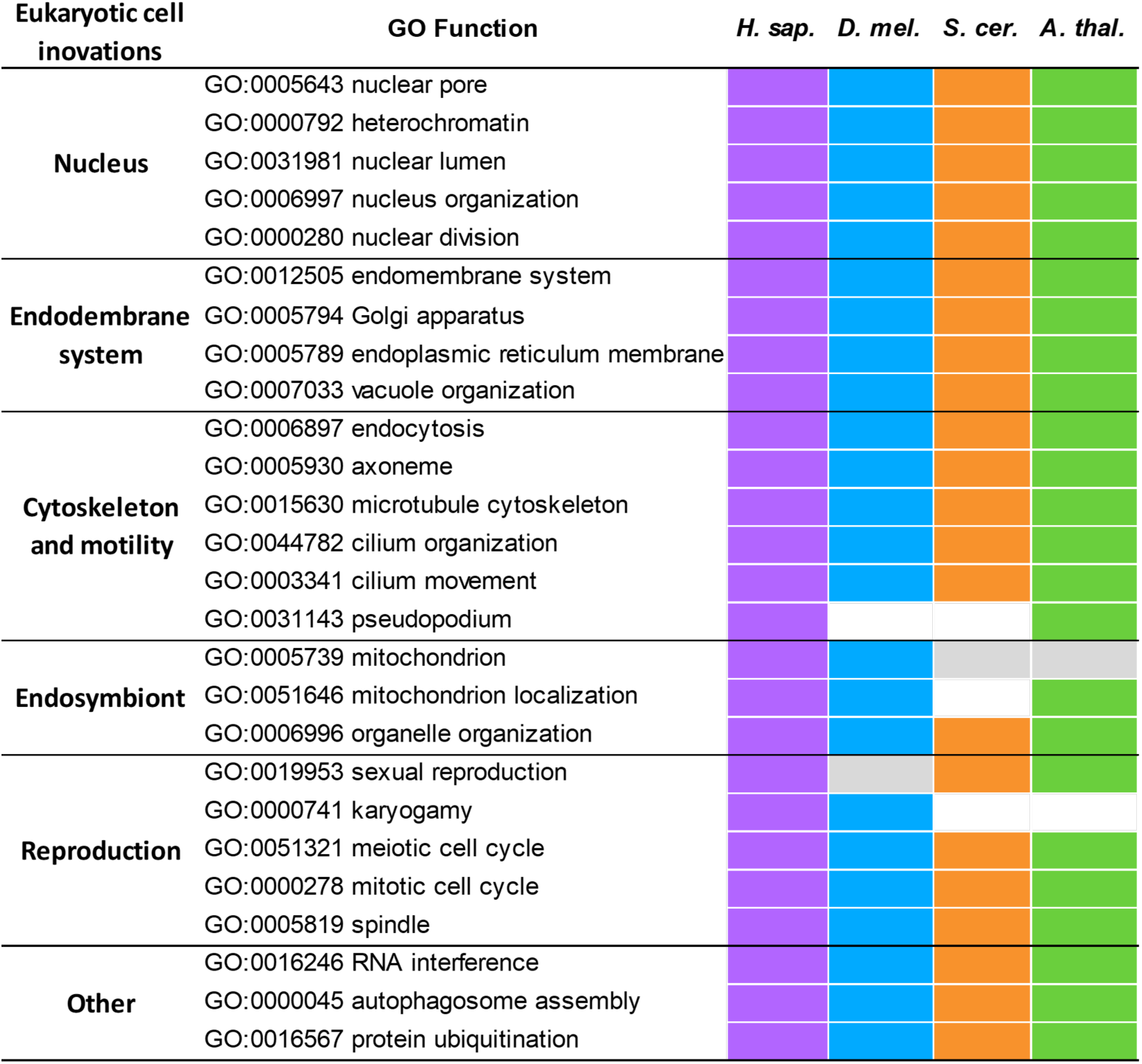
An overview of eukaryogenesis-related GO terms that show significant enrichment signals at Eukaryota (ps6). Colored fields mark the presence of the enrichment signal for a particular GO term at Eukaryota (ps6), gray fields mark that enrichment signal are present but not at ps6, and an empty field denotes complete absence of enrichment signals in a particular species. We performed an independent enrichment analysis for every focal species (*H. sapiens*, *D. melanogaster*, *S. cerevisiae*, and *A. thaliana*). The full enrichment profiles in the form of charts for each term and focal species are accessible in Supplementary File 15.

Beside these predictable patterns, we also uncovered some that are intuitive but previously undetected. An example is a strong enrichment of protein families functionally related to cilium/flagellum at the origin of Choanozoa (ps12), which are detectable from the perspective of *D. melanogaster* and *H. sapiens* as focal species (Fig. 6, 7). These strong enrichment signals at Choanozoa (ps12) partially overshadow those at the origin of Eukaryota (ps6), which are more prominent from the yeast and *Arabidopsis* perspective (Fig. 6,7). This pattern suggests that the flagellar apparatus of choanozoans, compared to other eukaryotes, is a derived feature underpinned by the recruitment of novel protein families.

**Figure 6.**
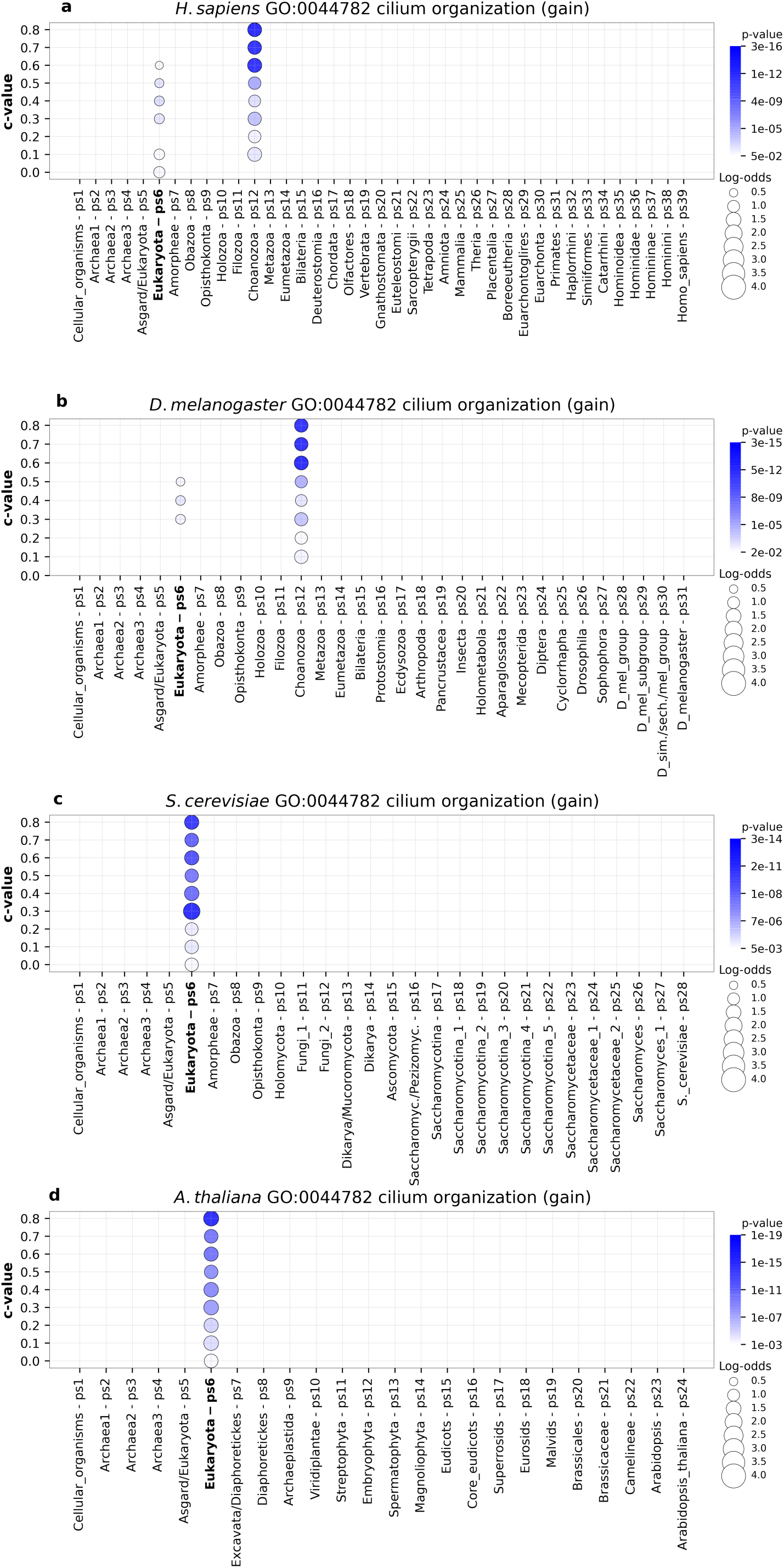
The enrichment of GO functional categories related to cilium organization in four focal species. The enrichment profiles are shown for the GO term GO:0044782 (cilium organization). The functional enrichments were calculated using the sets of gained gene families along **a**, *H.* sapiens, **b**, *D. melanogaster* **c**, *S. cerevisiae* and **d**, *A. thaliana* lineage (x-axis). The gene families are reconstructed with MMSeq2 using a range of c-values (0 to 0.8, y-axis). Solid circles depict significant enrichments of the GO term in gained gene families at a particular phylostratum. The size of circles is proportional to enrichment values estimated by log-odds, while the shades of blue (gain) correspond to p-values. The significance of enrichments was estimated by hypergeometric test corrected for multiple comparisons. Only enrichments with p < 0.05 are shown. This GO term shows the significant enrichments signals at the origin of Eukaryota (ps6) in all four focal species (**a-d**), with prominent additional signals at Choanozoa (ps12) in *H.* sapiens (**a**) and *D. melanogaster* (**b**) analysis.

**Figure 7.**
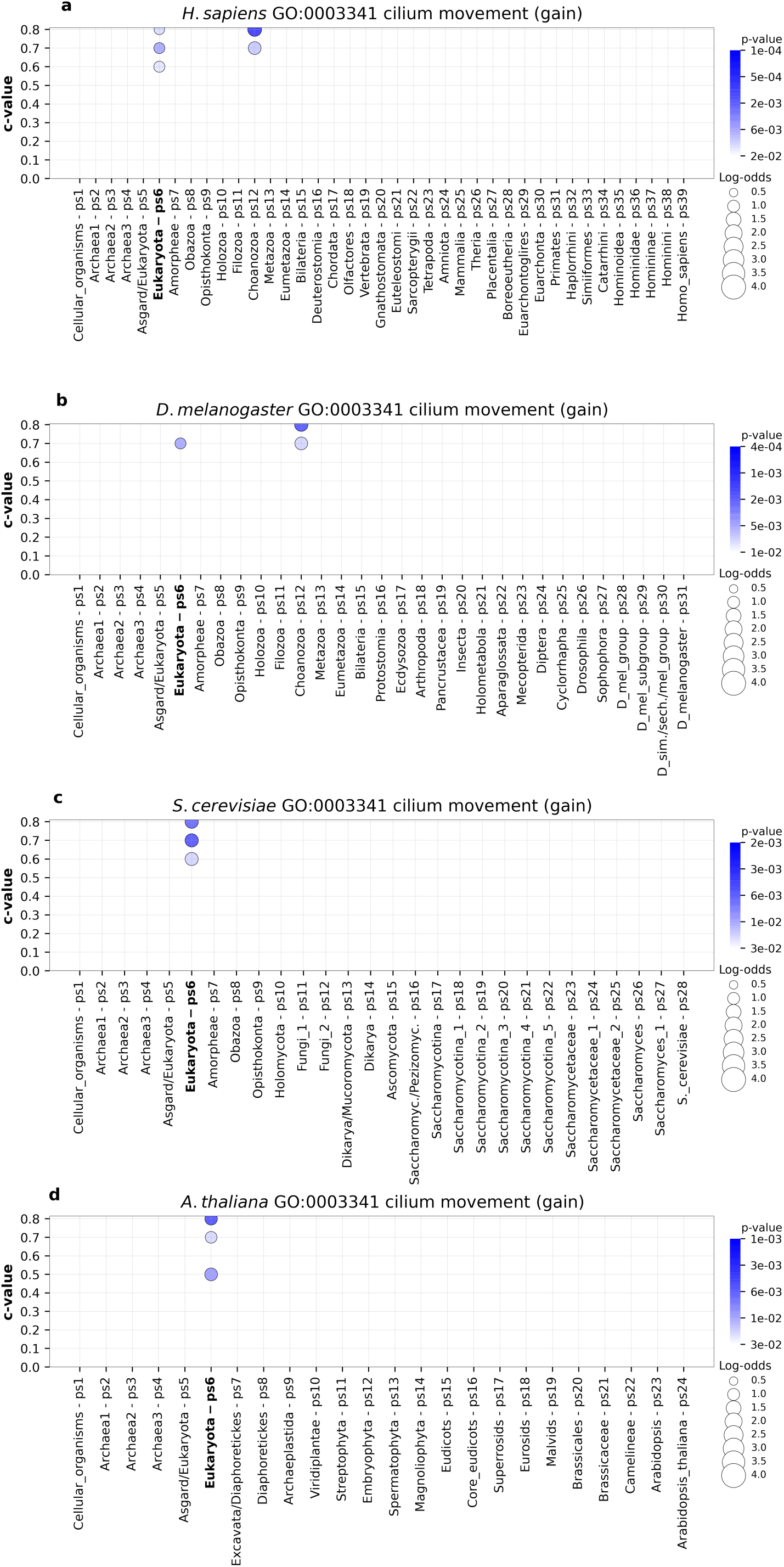
The enrichment of GO functional categories related to cilium movement in four focal species. The enrichment profiles are shown for the GO term GO:0003341 (cilium movement). The functional enrichments were calculated using the sets of gained gene families along **a**, *H.* sapiens, **b**, *D. melanogaster* **c**, *S. cerevisiae* and **d**, *A. thaliana* lineage (x-axis). The gene families are reconstructed with MMSeq2 using a range of c-values (0 to 0.8, y-axis). Solid circles depict significant enrichments of the GO term in gained gene families at a particular phylostratum. The size of circles is proportional to enrichment values estimated by log-odds, while the shades of blue (gain) correspond to p-values. The significance of enrichments was estimated by hypergeometric test corrected for multiple comparisons. Only enrichments with p < 0.05 are shown. This GO term shows the significant enrichments signals at higher c-values at the origin of Eukaryota (ps6) in all four focal species (**a-d**), with additional signals at Choanozoa (ps12) in *H.* sapiens (**a**) and *D. melanogaster* (**b**) analysis.

### Lineage specific GO functional enrichments

To further confirm the biological relevance of our gene family reconstruction, we explored GO functional enrichments that are specific for each of our four focal lineages. For instance, the emergence of adaptive immunity in vertebrates left a strong imprint in the evolutionary period that represents early vertebrate radiation (Fig. 8, Vertebrata to Placentalia, ps19-ps27), with especially prominent signals at the origin of jawed vertebrates (Gnathostomata, ps20). These functional enrichment patterns are in line with the view that the origin of adaptive immune system in vertebrates is a major evolutionary innovation (Müller et al. 2017, Flajnik et al. 2018). Similarly, metamorphosis is considered a key innovation that contributed to biological success of insects (Truman 2019). The enrichment profiles in *D. melanogaster* lineage reveal that the insect metamorphosis has deep roots in animal phylogeny and that it was continuously reshaped by the recruitment of novel protein families along the insect radiation (Fig. 9, Bilateria to Sophophora, ps15-ps28). Interestingly, we also detected significant loss of protein families related to metamorphosis in *D. melanogaster* (Fig. 9, ps31) in line with the previous findings of substantial gene family loss in this species (Hahn et al. 2007).

**Figure 8.**
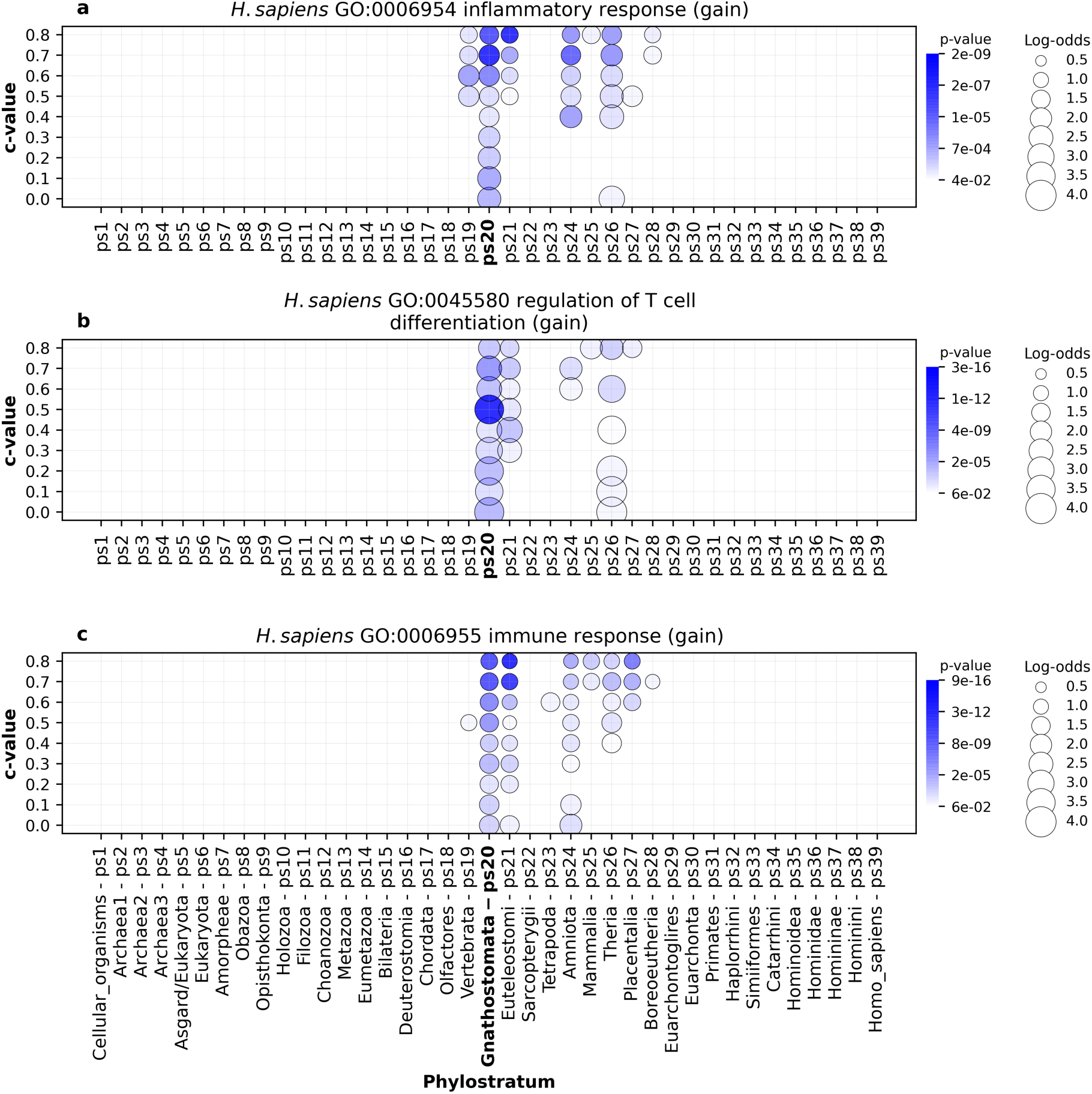
The enrichment of GO functional categories related to adaptive immune system (*H. sapiens*). The results are shown for three GO terms: **a**, GO:0006954 (inflammatory response) **b**, GO:0045580 (regulation of T cell differentiation), and **c**, GO:006955 (immune response). Functional enrichments were calculated using the set of gene families gained along *H. sapiens* lineage (x-axis). The gene families are reconstructed with MMSeq2 *cluster* using a range of c-values (0 to 0.8, y-axis). Solid circles depict significant enrichment of a GO term in gene families gained at a particular phylostratum. The size of circles is proportional to enrichment values estimated by log-odds, while the shades of blue correspond to p-values. The significance of enrichments was estimated by hypergeometric test corrected for multiple comparisons. Only enrichments with p < 0.05 are shown. These three terms show the strongest signal at the origin of jawed vertebrates (Gnathostomata, ps20), a group which evolved adaptive immune system. Other immune system related GO terms show similar patterns in *H. sapiens* lineage (Supplementary File 7).

**Figure 9.**
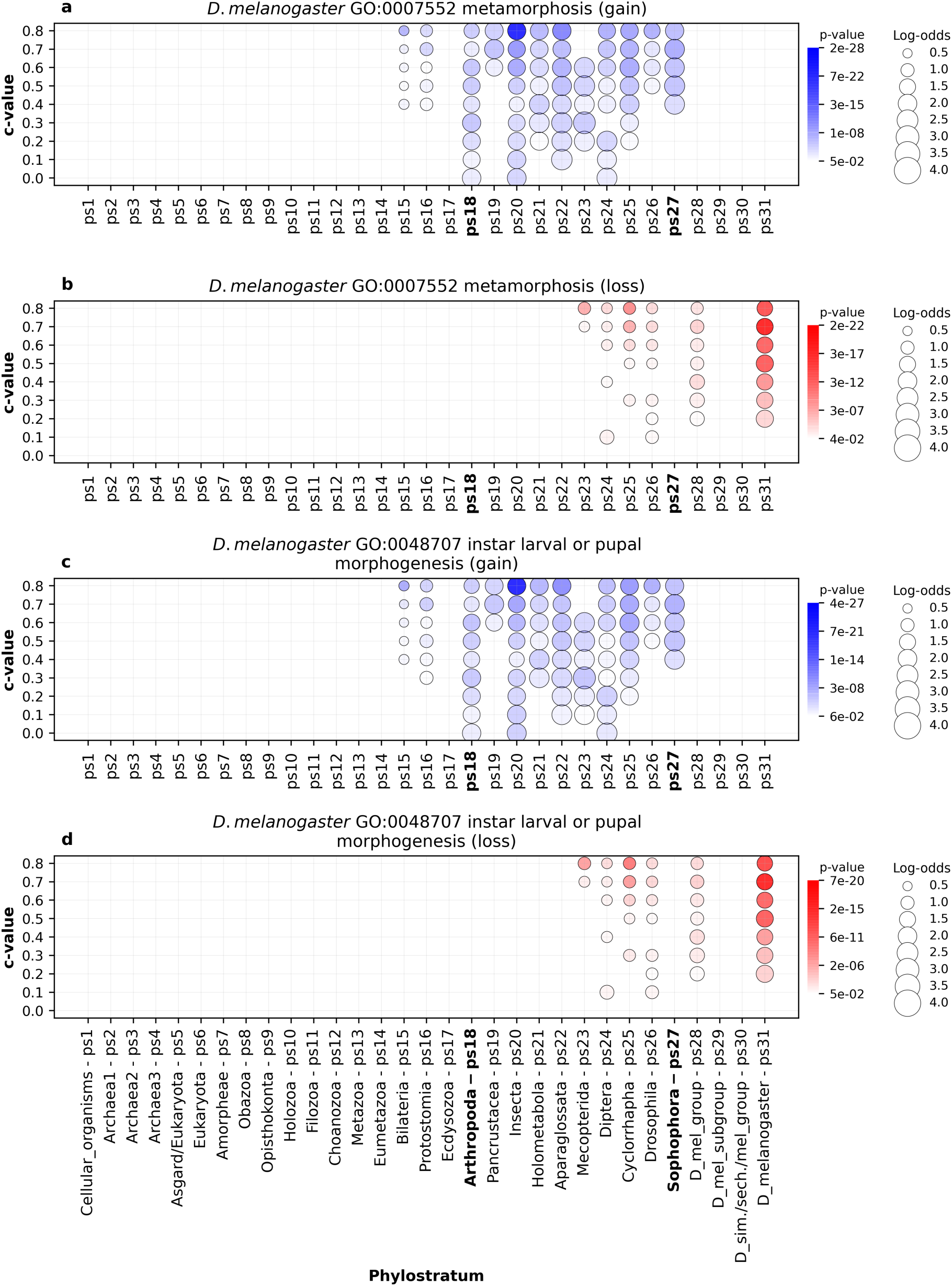
The enrichment of GO functional categories related to metamorphosis in insects (*D. melanogaster*). The enrichment profiles are shown for two GO terms: GO:0007552 (metamorphosis) and GO:0045580 (instar larval or pupal morphogenesis). The functional enrichments were calculated using the sets of gene families gained (**a, c**) and lost (**b, d**) along *D. melanogaster* lineage (x-axis). The gene families are reconstructed with MMSeq2 *cluster* using a range of c-values (0 to 0.8, y-axis). Solid circles depict significant enrichments of a GO term in gained or lost gene families at a particular phylostratum. The size of circles is proportional to enrichment values estimated by log-odds, while the shades of blue (gain) and red (loss) correspond to p-values. The significance of enrichments was estimated by hypergeometric test corrected for multiple comparisons. Only enrichments with p < 0.05 are shown. These two GO terms show the strongest gain-signals in the range between the origin Arthropoda (ps18) and the origin of Sophophora flies (ps27). Significant gene loss events are evident along diversification of dipterans (ps24-ps31).

A striking example of reductive evolution is the loss of respiratory chain complex I in the oxidative phosphorylation pathway of Saccharomycetaceae yeasts related to their fermentative and anaerobic lifestyles (Marcet-Houben et al. 2009, Schikora-Tamarit et al. 2021). This loss was initially detected by orthology tracking (Marcet-Houben et al. 2009); however, our protein family-based approach also recovered a statistically significant loss of functions associated to respiratory chain complex I at Saccharomycetaceae *sensu lato* (ps22, Fig. 10, Supplementary File 1).

**Figure 10.**
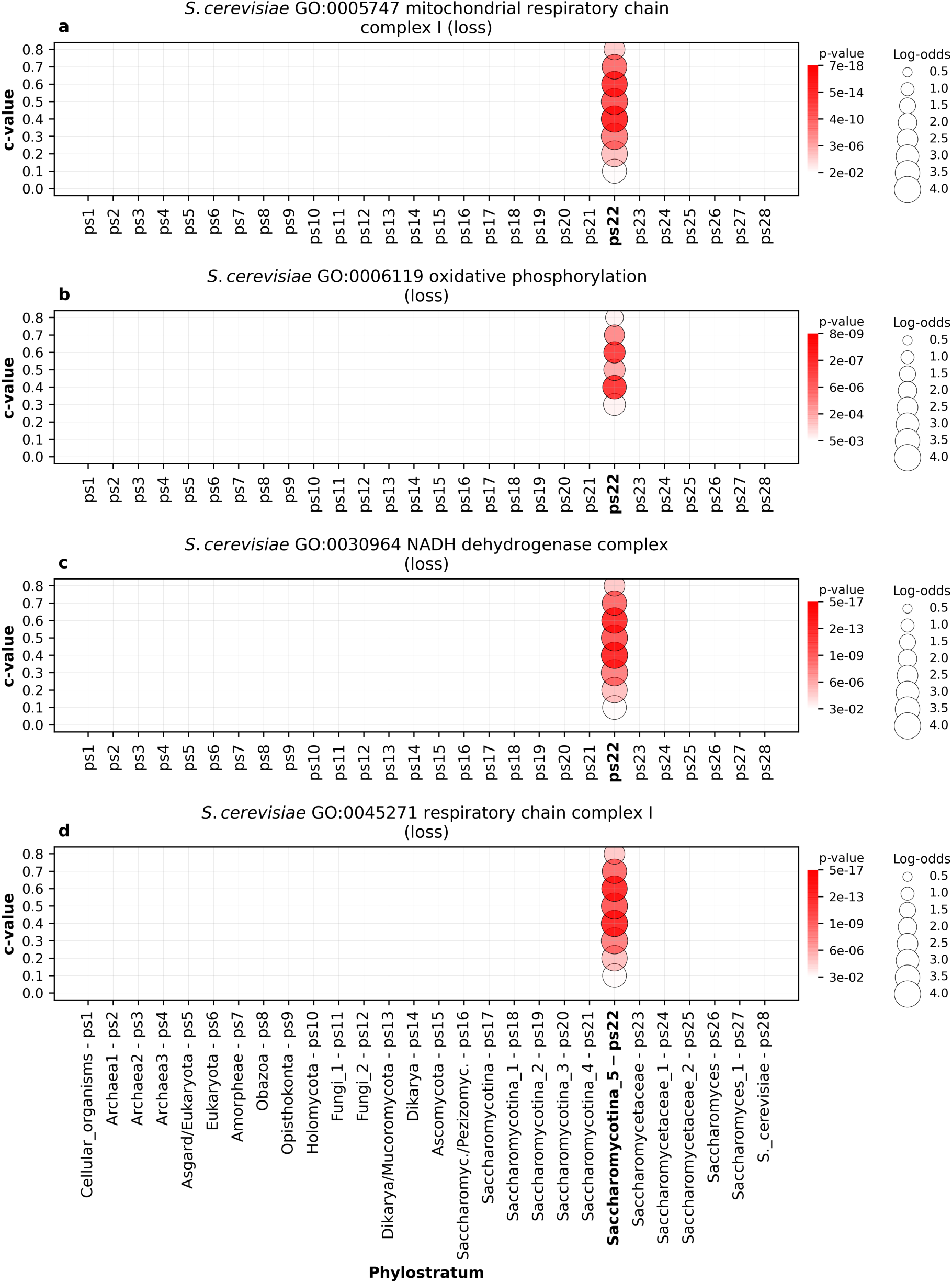
The enrichment of GO functional categories related to respiratory chain complex I (*S. cerevisiae*). The results are shown for four GO terms: **a**, GO:0005747 (mitochondrial respiratory chain complex I) **b**, GO:0006119 (oxidative phosphorylation), and **c**, GO:0030964 (NADH dehydrogenase complex) **d**, GO:0045271 (respiratory chain complex I). Functional enrichments were calculated using the set of gene families lost along *S. cerevisiae* lineage (x-axis). The gene families are reconstructed with MMSeq2 *cluster* using a range of c-values (0 to 0.8, y-axis). Solid circles depict significant enrichment of a GO term in gene families lost at a particular phylostratum. The size of circles is proportional to enrichment values estimated by log-odds, while the shades of red correspond to p-values. The significance of enrichments was estimated by hypergeometric test corrected for multiple comparisons. Only enrichments with p < 0.05 are shown. These four terms show the strongest loss signal at the origin of Saccharomycotina_5 (ps22), a phylostratum that represents Saccharomycetaceae yeasts (Supplementary File 1) that lost respiratory chain complex I.

Similar to other three focal species, we recovered the enrichment signals of many functions that mark the biology of *A. thaliana* lineage (Supplementary Files 1,10,14). Plastids, which originated by an endosymbiotic event in Archaeplastida, which involved cyanobacteria and eukaryotic host, are arguably the most important innovation that allowed radiation and ecological expansion of the plant lineage (de Vires et al. 2016). Our enrichment analysis uncovers continuous gain of protein families with plastid related functions in the broad period from the origin of Diaphoretickes to Eurosids (ps8-ps19, Fig. 11). The signal at Diaphoretickes (ps8, Fig. 11) reflects the phylogenetic distribution of plastids gained through secondary endosymbiosis (McGrath 2020), while signals in later periods suggest continuous and intensive coevolution of plastids and host cells in green algae (Viridiplanteae-Streptophyta ps9-ps10, Fig. 11) and along diversification of land plants (Embriophyta-Eurosids, ps11-ps19, Fig. 11). A prominent example of plastid evolution within angiosperms is emergence of chromoplasts, a derived plastid type, which confers bright colors to flowers and fruits (de Vires et al. 2016). In accordance with this, our analysis uncovered strong enrichment of gene families related to chromoplasts at the origin of flowering plants (ps14, Magnoliophyta, Fig. 11) and especially at core Eudicots (ps16, Fig. 11).

**Figure 11.**
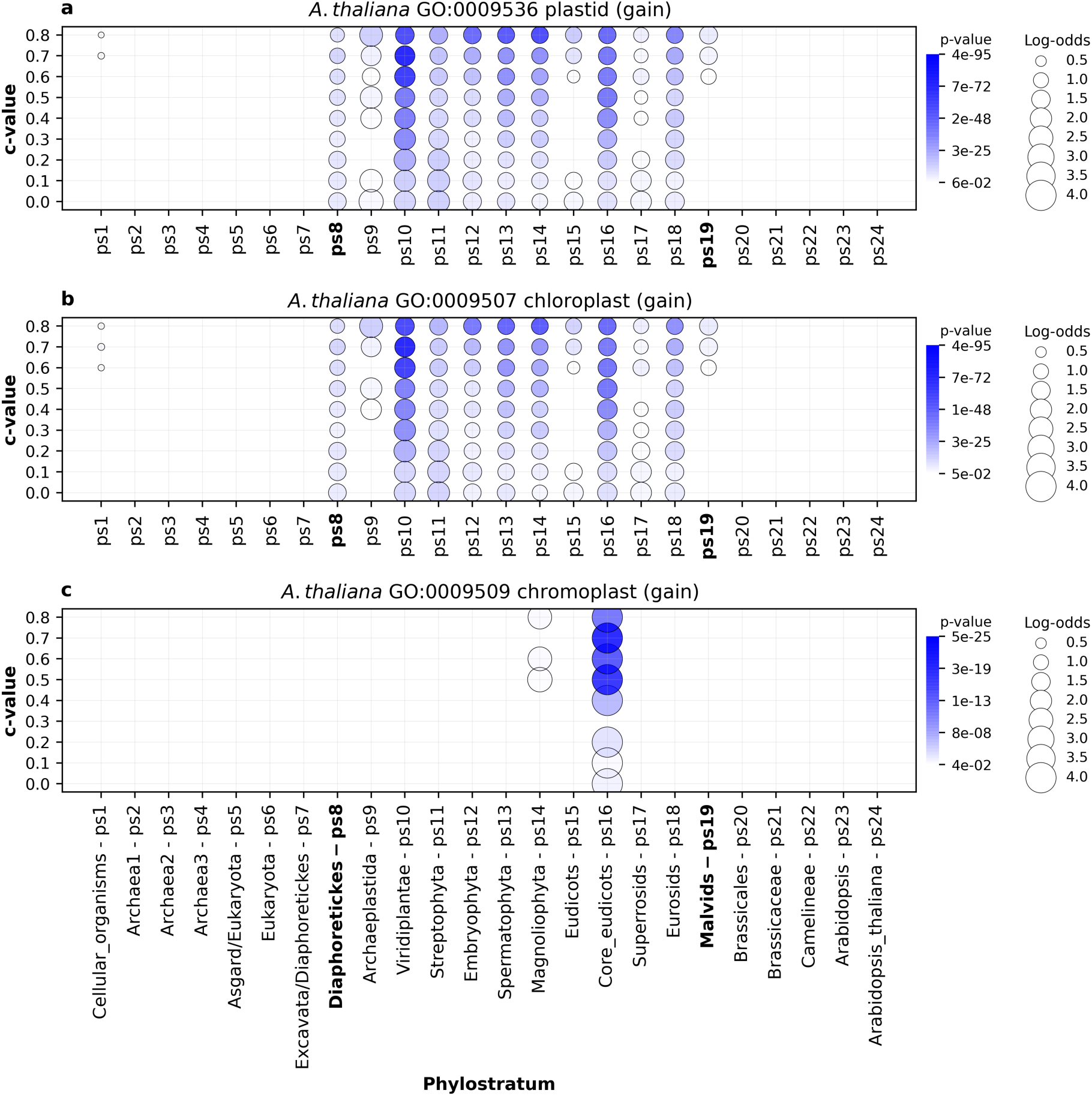
The enrichment of GO functional categories related to plastid evolution (*A. thaliana*). The results are shown for three GO terms: **a**, GO:0009536 (plastid) **b**, GO:0009507 (chloroplast), and **c**, GO:0009509 (chromoplast). Functional enrichments were calculated using the set of gene families gained along *A. thaliana* lineage (x-axis). The gene families are reconstructed with MMSeq2 *cluster* using a range of c-values (0 to 0.8, y-axis). Solid circles depict significant enrichment of a GO term in gene families gained at a particular phylostratum. The size of circles is proportional to enrichment values estimated by log-odds, while the shades of blue correspond to p-values. The significance of enrichments was estimated by hypergeometric test corrected for multiple comparisons. Only enrichments with p < 0.05 are shown. These three terms show the strongest loss signal in the range between Diaphoretickes and Eurosids (ps8-ps19). A week, but significant signal that reflects the origin of plasmids from cyanobacteria is evident at the origin of Cellular organisms (ps1, **a**, **b**).

Finally, the lineage specific profiles we described in more details here are only a tiny portion of those available in Supplementary Files 7-14, which could serve as a valuable resource for researchers interested in functional macroevolutionary patterns.

## Discussion

Taken together, the functional signals that we recovered in our gain-and-loss phylostratigraphic maps demonstrate biological validity of our gene family reconstruction, which shows the increase-peak-decrease pattern of protein family diversity at the macroevolutionary scale (Fig. 1). The shape of these protein diversity trajectories largely agrees with the predictions of biphasic and complexity-by-subtraction models of genome complexity evolution (Wolf and Koonin 2013, McShea and Hordijk 2013). However, it remains unclear which evolutionary forces produce these large-scale increase-peak-decrease protein diversity patterns that appear in independently evolving eukaryotic lineages. There is an extensive discussion on this topic in previous work that tries to weigh the relative importance of adaptive and neutral processes (Wolf and Koonin 2013, McShea and Hordijk 2013, O’Malley et al. 2016). The higher number of functional enrichments that we found in gene family gain compared to gene family loss events (Table 1) suggests that gene family losses are more frequently result of neutral processes. Yet, numerous functional enrichments we found in gene family loss patterns indicate (Supplementary Files 11-14) that many reductive events are the result of adaptive evolution. For example, the loss of respiratory chain complex I in yeasts (Fig. 10) is likely an adaptive event, although at present it can be only speculated which selective pressures drove this reduction (Marcet-Houben et al. 2009, Schikora-Tamarit et al. 2021, O’Malley et al. 2016).

Nevertheless, on the gross scale it is very indicative that the reductive trend in our focal lineages correlate with important ecological transitions. For instance, after the split of deuterostomic and protostomic animals the complexity of their genomes at the level of gene families were continuously reduced until the present time (Fig. 1a,b). This switch to the reductive mode of genome evolution corresponds to Proterozoic-Cambrian transition, where profound abiotic and biotic environmental changes occurred, which, in turn, allowed animal radiation and the complexification of marine ecosystems (Sperling et al. 2013, Wood et al. 2019, Zhang et al. 2021). One could speculate that the rise in complexity of feeding ecology in this geological period, which included the evolution of more efficient predation modes, together with the emergence of new and more specialized ecological niches (Sperling et al. 2013, Wood et al. 2019, Zhang et al. 2021), created possibilities for genome reduction.

A support for this idea comes from the free-living communities of planktonic bacteria where some members provide costly and indispensable functions as public goods. Other bacterial species in the community exploit the environmental availability of these functions and benefit by simplifying their genomes via the loss of costly genetic machineries (Morris et al. 2012). The Black Queen Hypothesis (BQH), which models this phenomenon, predicts that the loss of such leaky and costly functions is selectively favored, and that it leads to the emergence of new and long-lasting biological dependencies between bacterial species (Morris et al. 2012). This idea could be generalized in a way that any function that is costly to perform for an organism and at the same time it could be outsourced through biological interactions is a potential target of reductive evolution. These processes could leave then imprint in the genome by decreasing gene family diversity of an organism. We term this generalization “functional outsourcing”. For instance, changes in feeding ecology, niche specialization, increasingly more intimate interactions within animal holobionts (McFall-Ngai et al. 2013), could all have triggered the reduction in protein family diversity within particular animal lineages.

Interestingly, if this idea holds true, our patterns of continuous reduction in protein family diversity during Phanerozoic period (Fig. 1a,b) signal that the complexity and strength of biological interactions, which involve animals, were more or less continuously increasing in the last 540 million years. In other words, gene family diversity (complexity) is negatively correlated with the strength (complexity) of biological interactions. This sheds some light on the problem of measuring complexity of biological systems. It is well known that organismal complexity, at various phenotypic levels, often does not reflect genomic complexity, and vice versa (Wolf and Koonin 2013). Wolf and Koonin proposed that genomic complexity measured by the number of conserved genes could be complemented with other measures to obtain a better proxy of organismal complexity (Wolf and Koonin 2013). To our knowledge there have not been any further attempts in this direction so far.

To contribute to this idea, we here propose that complexity of an organism might be estimated by the number of conserved functions that are hardcoded in its genome (e.g. gene families) plus the number of functional benefits achieved through the direct biological interactions with other organisms (e.g. gene families in interacting organisms that contribute to that benefit). This way of viewing organism complexity goes beyond the holobiont paradigm (McFall-Ngai et al. 2013) and moves the focus from the host and its microbes to the complete ecological community that interacts with a particular organism. In this respect it would be important to distinguish between functions that are lost and are not needed any more, and those that are lost but outsourced, as only the latter contribute to organismal complexity. For instance, the loss of essential amino acids synthesis capability in metazoans (Richter et al. 2018) is compensated by the digestion of other organisms or through gut microbiota symbiosis. In this case complexity score would not change, because other organisms produce essential amino acids for metazoans which harness them through biological interaction. However, an opposite example would be the loss of cilia in fungi (Naranjo-Ortiz and Gabaldón 2019) because this functionality is not outsourced; the function is simply lost and not needed anymore in the fungal lifestyle.

If one extends this reasoning further to fungi, then very strong reductive trend in protein family diversity, which starts already at Holomycota (ps10, Fig. 1c), suggests that fungi very early adopted lifestyles that include biological interactions, which allowed them to reduce the pack of protein families necessary for survival. Indeed, fungi evolved most likely from predatory protists that switched to parasitism (Naranjo-Ortiz and Gabaldón 2019). Later, in the course of their evolution they diversified by making numerous ecological transitions to predatory, pathogenic, parasitic, and symbiotic interactions (Naranjo-Ortiz and Gabaldón 2019), all of which allow ecological specializations and lower the need for self-production of many protein families.

However, the pattern in plants is more intricate (Fig. 1d). After eukaryogenesis was completed (ps6, Fig. 1d), there is an obvious simplification trend in the lineage leading to plants with the lowest values at the origin of Streptophyta (ps11, Fig. 1d). This reduction in gene family diversity suggests increasing ecological complexification of aquatic ecosystems where these organisms thrived (de Vires et al. 2016, de Vires and Archibald 2018). Nevertheless, after the colonization of land begun with Embryophyta (ps12, Fig. 1d), an opposite trend of increase in gene family diversity started (ps12, Fig. 1d), which finally reached the peak at the origin of Eurosids (ps18, Fig. 1d). This increase probably reflects new adaptations, via adaptive recruitment of new gene families, to harsh conditions that plants had to face during challenging transition to land (Maberly 2014, de Vires and Archibald 2018). Finally, along the diversification of Eurosids (ps19-ps24, Fig. 1d) a strong simplification trend is evident, which suggests increase in biological interactions that allowed gene family diversity reduction. Indeed, rapid diversification of Rosids that started in Cretaceous period resulted in formation of rosid-dominated angiosperm forests present today (Wang et al. 2009). This rosid radiation included nutrient and habitat specializations as well as coevolution with animals, especially insects and mammals (Wang et al. 2009). All of this could create conditions for gene family diversity reduction. A striking example in angiosperms that demonstrates the impact of nutrient and habitat specializations on the genome content are carnivorous plants, whose genomes show massive gene loss connected to functional outsourcing (Palfalvi et al. 2020, Nevill et al. 2019).

From the technical side, our analysis also uncovered that a minimal alignment length, as controlled by c-value, is an important parameter that modulates amount of macroevolutionary signal that could be recovered on the gene family gain-and-loss phylostratigraphic maps. In general, higher c-values that impose stringent criteria on the sequence architecture carry more macroevolutionary information (Supplementary Table 2), although in some instances lower c-values provide evolutionary informative patterns that are not visible at higher c-values (e.g., ps6 Fig. 6a,b). We thus propose that, instead of choosing one cut-off value, the best strategy is to explore protein sequence space using a broad range of c-values and then inclusively evaluate evolutionary signals at hand.

Our result that higher c-values typically carry more biological information is also relevant in the context of debate on the importance of deep homologs for macroevolutionary reconstruction and the ability of sequence similarity search algorithms to detect them (Moyers and Zhang 2015, Domazet-Lošo et al. 2017, Natsidis et al. 2021, Futo et al. 2021). Our study clearly showed that functional information recovered by the enrichment analysis increasingly erodes in remote homologs (Supplementary Table 2, Supplementary Files 7-14), which makes them less useful in tracking functional evolution. However, studies that use sequence divergence simulations in an attempt to challenge macroevolutionary patterns obtained using real datasets, do not consider how the sequence divergence of artificially evolved sequences translates to their functional divergence (Moyers and Zhang 2015, Natsidis et al. 2021). This brings into question the biological relevance of these simulation studies, and probably explains why the effort to link simulated sequences with functional evolutionary patterns failed (Moyers and Zhang 2015, Moyers and Zhang 2016, Domazet-Lošo et al. 2017). Finally, if we consider all genes that evolved by perpetual duplication processes across the tree of life, then essentially all of them are deep homologs that coalesce, irrespective of their sequence similarity or lack of it, to some primordial sequence (Domazet-Loso and Tautz 2003). The grouping of all these genes as homologs to the oldest phylogenetic node, which is a default expectation in the current simulation studies (Moyers and Zhang 2015, Natsidis et al. 2021), obviously ignores the plethora of selective pressures that shaped their sequence divergence over macroevolutionary time. In consequence, this procedure is functionally completely uninformative.

Another important aspect of this work relates to the benchmarking of computationally recovered gene family clusters in terms of their evolutionary and biological validity. There is a general consensus on the evolutionary origin of some biological features like eukaryogenesis related innovations, origin of the adaptive immune system in animals, plastid endosymbiosis or the loss of respiratory chain complex I in yeasts. These and similar features could then be used to evaluate and calibrate bioinformatic pipelines with an aim to maximize the biological information that could be recovered by statistical analysis. This will also allow us to contrast the biological information content in gene clusters generated via different types of grouping strategies (e.g., orthologs vs homologs).

In conclusion, our study demonstrates that the patterns of gene family gain and loss correlate with major evolutionary and ecological transitions. It seems that the gene family loss follows the evolutionary radiations of major multicellular eukaryotic lineages as a consequence of ecological complexification that allowed niche specializations and functional outsourcing. This in turn suggests that in evaluating the complexity of an organism, in addition to the number of its conserved parts (e.g., gene families), one should also consider its biological interactions and the functional context that sustain its existence.

## Methods

### Consensus phylogeny

Using information from the relevant phylogenetic literature, e.g. (Irisarri et al. 2017, Misof et al. 2014, Shen et al. 2016, Morris et al. 2018), we constructed a consensus tree with 667 species taxa, whose genomes were publicly available (Supplementary File 1). In the assembly of the tree, we aimed to comprehensively cover lineages that lead from the origin of cellular organism to our four focal species; i.e., *H. sapiens*, *D. melanogaster*, *S. cerevisiae* and *A. thaliana*. We chose these focal species because they are well-studied model organisms, which allowed us to reach an adequate number of phylogenetic levels (phylostrata) and to populate them with suitable amount of reference genomes. In addition, these lineages represent the evolutionary trajectories of deuterostomic animals, protostomic animals, fungi and plants, all of which are ecologically important and evolutionary successful multicellular eukaryotic groups with currently the best annotation of gene functions.

### Reference genomes

We retrieved the full protein sequence sets (reference genomes) of our 667 taxa from the ENSEMBL and NCBI databases. In addition, we retrieved some reference genomes, which were not available in these databases, from other sources (Supplementary Table 3). In every reference genome, we retained only protein sequences that come from the longest splicing variant of a respective gene. To detect possible low-quality reference genomes, we tested their completeness using BUSCO v4.04 with default parameters by performing analysis with eukaryota_odb10, bacteria_odb10 and archea_odb10 datasets (Mani et al. 2021). We then individually assessed every genome and if the quality looked compromised, we substituted it with a more complete version, or alternatively with a better-quality genome of a closely related species. The list of reference genomes and their corresponding BUSCO scores are available in Supplementary Table 3.

### Sequence similarity search and clustering

The protein sequences of all reference genomes were clustered using *mmseqs cluster* algorithm within the MMseqs2 package (Hauser et al. 2016, Steinegger and Söding 2017). This algorithm integrates all-against-all protein sequence similarity search with clustering procedure. We retained *mmseqs cluster* default e-value cutoff of 10^-3^ as this threshold was repeatedly shown to be optimal (Domazet-Loso and Tautz 2003; Domazet-Lošo et al. 2017; Vakirlis et al. 2020, Futo et al. 2021). To cluster our set of protein sequences, we applied connected component algorithm (*--cluster-mode 1*). In contrast to other available clustering options in *mmseqs cluster* (Steinegger and Söding 2018), this is a transitivity-based clustering algorithm that forms clusters with more remote homologs. We independently clustered our protein sequence set nine times by varying c-values in the range between 0 and 0.8 with 0.1 increments. A c-value determines the minimal percentage of sequence length (0% to 80%) that is required for the assignment of a protein sequence to a cluster. The alignment coverage was calculated under *--cov-mode 0* which calculates the percentage of alignment overlap in reference to a longer protein sequence. The higher percentages of alignment coverage ensure that all sequences within a cluster share increasingly similar overall sequence architecture (stringent criteria). In contrast, lower c-values permit clustering based on the conservation of shorter protein sequence stretches irrespective of their arrangement within genes (permissive criteria). This approach allowed us to explore evolutionary relevant shifts in the protein sequence space at the level of short conserved protein sequence stretches (c-value 0), at one extreme, and at the level of sequence conservation along almost complete gene length (c-value 0.8) at the other one.

In contrast to previous studies that rely on orthology-focused clustering that filters out a part of sequence similarity information (Richter et al. 2018, Guijarro-Clarke et al. 2020, Fernández and Gabaldón 2020), e.g. by considering only the reciprocal best hits (Paps and Holland 2018, Guijarro-Clarke et al. 2020), our clustering protocol takes into account all detectable sequence similarity below an e-value cut-off and above a c-value threshold to form clusters. This approach allowed us to unrestrictedly explore protein sequence space and determine significant shifts in the form of gene family gains and losses. We repeatedly showed previously that these detectable jumps in the protein sequence space carry biological information, which can then be statistically recovered (Domazet-Lošo and Tautz 2003, Domazet-Lošo et al. 2007, Domazet-Lošo and Tautz 2010a, Domazet-Lošo and Tauz 2010b, Domazet-Lošo et al. 2017, Futo et al. 2021).

### Gene family gain and loss reconstruction

The taxonomic composition of every cluster (gene family) obtained by *mmseqs cluster* was mapped on the consensus tree. We then extracted only those gene families that appear in the lineages that lead from the root of the tree to one of our four focal species. In this way, each focal species obtained a set of gene families relevant for its lineage. We stress that every of the 667 species on the consensus phylogeny could be treated as a focal species and for each of them gene families relevant for their lineages could be extracted using the scripts we provide. However, this would create an overwhelming amount of redundant data that at present would be hard to put in the comparative context, therefore we restricted our analysis to the four most important representative species. By applying Dollo’s parsimony, we then determined for every gene family the phylostratum of its gain and eventually of its loss along the focal-species lineage. Using this gene family gain-and-loss information we calculated gene family content in every ancestral node along a focal lineage (Supplementary Table 1, Fig. 1). It is important to note that a gene family gain event must always precede its loss by at least one internode (i.e., two phylostrata). This simply reflects the fact that a gain of a gene family in a particular phylostratum and its loss immediately in the next one excludes that gene family from the focal lineage and makes it specific for the respective side branch. For this reason, the loss events in a particular focal lineage could be calculated the earliest at the third phylostratum from the root.

### Functional enrichment analysis

We functionally annotated protein sequences in the reference genomes with COG and Gene Ontology (COG) annotations using *emapper* version 1.0.3-5, the database version 4.5.1 and *diamond* search algorithm within the EggNOG tool (Huerta-Cepas et al. 2019). To functionally annotate clusters (gene families) with either COG or GO terms, we applied a simple rule that every functional annotation of a protein member within a cluster is also assigned to the whole cluster. To calculate a COG or GO term functional enrichments in gained or lost gene family sets along focal species lineages, we performed one-tailed hypergeometric test for each focal species and c-value independently. To correct for multiple testing, we adjusted p-values using Benjamini–Hochberg method as implemented in the Python *statsmodels* library.

## Data availability

Publicly available genomes are listed in Supplementary Table 3. The scripts used in this study are available in GitHub at https://github.com/PhyLoss/PhyLoss.

## Acknowledgments

We thank M. Futo, S. Koska, N. Čorak and D. Kifer for discussions. This work was supported by the Croatian Science Foundation under the project IP-2016-06-5924, the City of Zagreb, the Adris Foundation and the European Regional Development Fund (KK.01.1.1.01.0009 DATACROSS). We used the computational resources of the University Computing Center - SRCE (Isabella) and the Ruđer Bošković Institute.

## Author contribution

T.D.-L. initiated the study, T.D.-L. and M.D.-L. conceptualized the study and constructed the phylogeny. M.D.-L. prepared the genomic data, developed the phylostratigraphic pipeline and the algorithm for the computation of gene family gain and loss. T. Š. performed functional annotations of clusters. T. Š. and M.D.-L. wrote the scripts for the functional analysis. All authors analyzed the data. M.D.-L. prepared the figures and tables for publication. M.D.-L. and T.D.-L. wrote the manuscript. All authors read and approved the manuscript.

